# Dynamic updating of hippocampal object representations reflects new conceptual knowledge

**DOI:** 10.1101/071118

**Authors:** Michael L. Mack, Bradley C. Love, Alison R. Preston

## Abstract

Concepts organize the relationship among individual stimuli or events by highlighting shared features. Often, new goals require updating conceptual knowledge to reflect relationships based on different goal-relevant features. Here, our aim is to determine how hippocampal (HPC) object representations are organized and updated to reflect changing conceptual knowledge. Participants learned two classification tasks in which successful learning required attention to different stimulus features, thus providing a means to index how representations of individual stimuli are reorganized according to changing task goals. We used a computational learning model to capture how people attended to goal-relevant features and organized object representations based on those features during learning. Using representational similarity analyses of functional magnetic resonance imaging data, we demonstrate that neural representations in left anterior HPC correspond with model predictions of concept organization. Moreover, we show that during early learning, when concept updating is most consequential, HPC is functionally coupled with prefrontal regions. Based on these findings, we propose that when task goals change, object representations in HPC can be organized in new ways, resulting in updated concepts that highlight the features most critical to the new goal.

**Significance Statement:** A cosmopolitan couple looking for a home may focus on trendy features. But, with news of a baby on the way, they must quickly learn which features make for a child-friendly home to conceptually reorganize their set of potential homes. Here, we investigate how conceptual knowledge is updated in the brain when goals change and attention shifts to new information. By combining fMRI with computational modeling, we find that object representations in the human hippocampus are dynamically updated with concept-relevant information during learning. We also demonstrate that when concept updating is most consequential, the hippocampus is functionally coupled with neocortex. Our findings suggest that the brain reorganizes when concepts change and provide support for a neurocomputational theory of concept formation.

---

Concepts are organizing principles that define how items or events are similar to one another. Goals are critical to shaping concepts, by emphasizing some shared features over others. When goals change, previously experienced events may be organized in new ways, resulting in an updated concept that highlights the features most critical to the new goal. For instance, consider purchasing a home. One must learn which features make for the most desirable home. A young couple seeking a cosmopolitan lifestyle may organize potential houses based on trendy features like exposed brick walls, a wet bar, and room for vintage record collections. However, with the news of a baby on the way, the couple’s goals are likely to shift. After pouring through parenting books and web forums to learn what makes for a child-friendly home, they may look at those previously seen potential homes in a different light. Instead, family-oriented features such as whether or not a home has a bathtub, is within walking distance to a park, and is in a well-respected school district may matter more resulting in a reorganization of which homes are a good buy. At the core of this example are the fundamental challenges we face in flexible goal-directed learning. When learning new concepts (e.g., child-friendly instead of a trendy house), attention changes focus to different information and items that were conceptually dissimilar (e.g., two houses with and without a wet bar) may become more similar (e.g., they both are close to a park) and vice versa (1). Understanding how conceptual knowledge is created and updated during learning is a central question for both cognitive psychology (1–3) and neuroscience (4–7); yet, few studies attempt to bridge these domains. Here, we test a neurocomputational account of concept formation by combining human functional magnetic resonance imaging (fMRI) with a computational model of learning.

We evaluate the proposal that during new learning, concept-relevant features are preferentially encoded into object representations in the hippocampus (HPC). Recent findings suggest HPC plays an important role in forming representations that integrate across shared features of experiences (6, 8–10); yet there is little understanding about how HPC representations evolve when conceptual knowledge changes. Prominent computational theories posit that concept formation in HPC is influenced by selective attention mechanisms that favor goal-relevant features from our experiences (1, 11, 12). When new goals arise, conceptual coding in HPC is reorganized according to the newly-relevant features selected by attention. Two lines of empirical evidence support this theoretical view. First, HPC rapidly learns (13), an ability important for updating conceptual representations in the face of changing goals. Second, HPC has also been shown to activate representations that are goal relevant (14–19). A critical open question is how the *same* experiences come to be represented differently in neural terms as a function of changing conceptual knowledge. We test the hypothesis that HPC coding, in concert with selective attention, builds and updates concepts, resulting in distinct representations for the same stimuli across different learning contexts.

Participants were first exposed to images of insects, which had three varying features (Fig. 1a). During high-resolution functional magnetic resonance imaging (fMRI) scans, participants learned two categorization problems using the insect stimuli. For one categorization problem (referred to as type 1 (20, 21)), participants learned to group the insects based on a single feature. For instance, participants were asked to sort insects into those that prefer warm or cool environments. Via trial and error, participants learned that an insect’s preference could be determined by attending to the width of the legs, with thick-legged insects preferring warm environments and thin-legged insects preferring cool environments. The other categorization program (termed type 2) required participants attend to the other two features (e.g., antennae and pincers) to perform correctly. For this problem, participants might be asked to sort the insects according to the hemisphere in which they are typically found, Eastern or Western. The correct conceptual grouping takes the form of an XOR rule; Eastern hemisphere insects comprised the insects with thick antennae and scooped pincers or thin antennae and sharp pincers, whereas Western insects were those with thick antennae and sharp pincers or thin antennae and scooped pincers. The order in which participants experienced these tasks was counterbalanced; half of participants learned the type 1 problem first, and the remaining participants learned type 2 first. Thus, the same stimuli were used in both learning tasks, but the conceptual mappings of the stimuli changed across tasks. To perform efficiently, participants had to learn to attend to different features of the insects and update their concepts when the task, and therefore the goal, changed (Fig. 1b).

**Figure 1:**
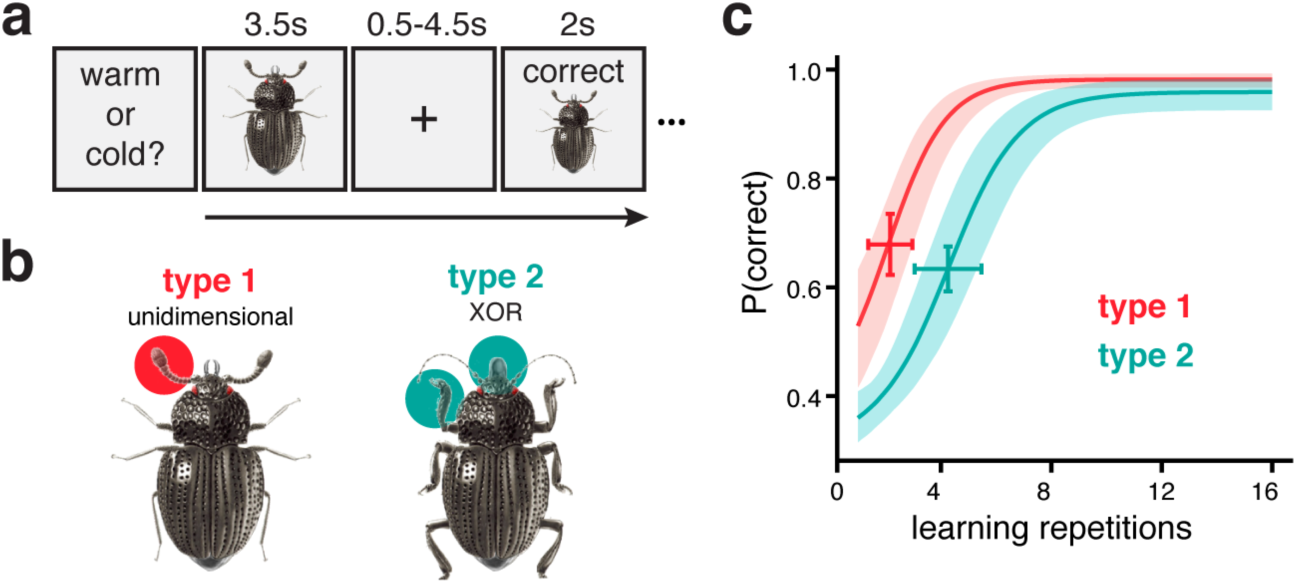
Experiment schematic and behavioral performance. **a)** Participants learned to classify eight insect images according to two different rules through feedback-based learning. On every trial, an insect image was presented (3.5s) and participants made classification responses according to the current task. After a delay (0.5-4.5s), feedback consisting of the insect image, the accuracy of the response, and the correct classification was shown (2s). The next trial began after a variable delay (4-8s). For both tasks, participants responded to all eight stimuli over sixteen repetitions. **b)** The stimuli consisted of insects with three binary features (thick/thin legs, thick/thin antennae, and pincer/shovel mouths). The stimulus set consisted of eight images representing all combinations of the three binary features. The two classification tasks required attention to different features: the type 1 problem was based on one feature (e.g., the antennae), the type 2 problem was an XOR classification based on a combination of two features (e.g., the mouth and legs). The feature-to-task mappings and order of the learning tasks were counterbalanced across participants. **c)** The average probability of a correct response across the 16 learning repetitions is plotted for both tasks. Error bars represent 95% confidence intervals (CI) around the inflection point of the bounded logistic learning curves. The shaded ribbons represent 95% CI of the mean.

This manipulation thus allowed us to vary the relevancy of the stimulus dimensions over time. By holding the stimuli constant and varying which features should be attended to across tasks, the features that were once relevant become irrelevant and the items that were once conceptually similar may become very different. For example, two insects that were considered similar in the first task because they share thin legs may become conceptually dissimilar in the second task because they have different antennae or mouths. The change in feature relevancy therefore requires rapid updating of conceptual representations, both initially after the exposure phase and in the transition from one task to another. Using a computational learning model named SUSTAIN (1), we created formal predictions about how concepts were updated for each task. This learning model (Fig. 2a) is based on two central mechanisms: 1) attention weights to stimulus features and 2) conceptual knowledge stores, called clusters, that represent weighted combinations of feature values and an association to a class label. A classification decision is made by first weighting stimulus feature values according to the attention weights, then comparing the attention-weighted stimulus inputs to the stored clusters. The most similar cluster is then used to drive a probabilistic decision. Importantly, this model predicts learning behavior through a feedback-driven process that tunes the attention weights to select features most informative for the current task. The clusters are also adaptively updated to code for the similarities among the stimuli that best represent the concepts needed for the current task. In other words, the model optimizes the organization of cluster representations over the course of learning based on changing task goals and the stimulus features that are most task relevant.

**Figure 2:**
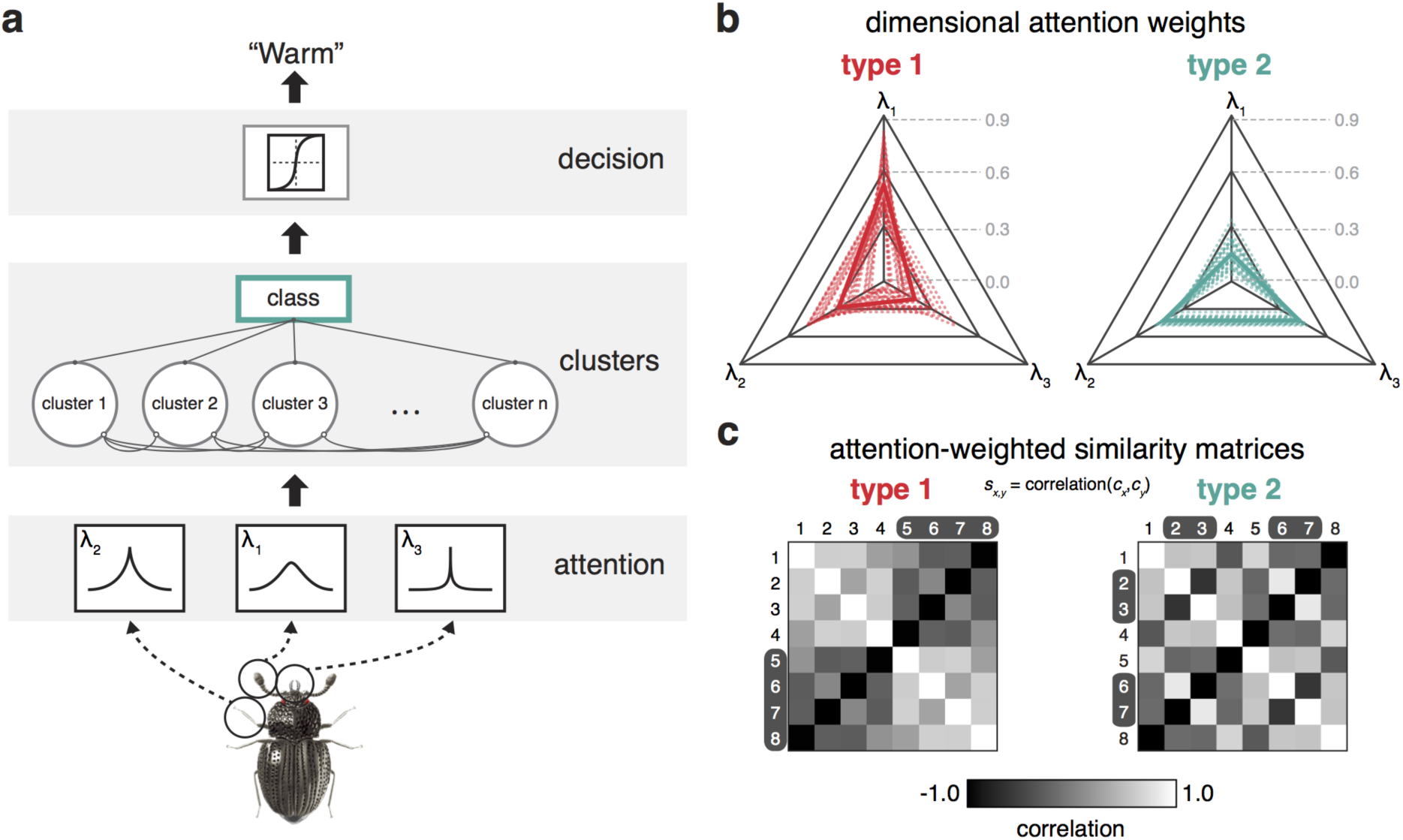
Schematic of learning model and model predictions. **a)** The learning model consists of three main components (see Supporting Information for model formalism). First, the sensory input of the three stimulus features is attenuated by receptive field filters tuned according to attention weights (*λ*_i_). The attention component acts to alter the perceptual representation of the stimulus towards task-diagnostic information. Second, stored knowledge represented by clusters of weighted features compete to be activated by the attention-biased input. The cluster most similar to the attention-biased input wins and activates the class unit. Third, the activated class unit serves as input to a decision component that generates a response. Trial-to-trial, the model learns through feedback by updating the attention weights and the weights connecting clusters to the class unit, and whether an existing cluster is updated or a new cluster is recruited. **b)** The model was fit to participants learning performance (Fig. 1c) and the final attention weights (*λ*_i_) for each dimension were extracted for both tasks. The relative attention weights for each task are depicted in the radar plots (dotted lines show participant weights, bold lines show group means). **c)** Matrices depict the average model predictions for the pairwise similarities between the stimuli for the two tasks. Task-specific similarity predictions for each participant were generated by extracting cluster activations for each stimulus at the end of learning. Pearson correlations were then calculated for each stimulus pair, and averaged across participants. The similarity matrices characterize the task-specific conceptual representations underlying classification decisions. Stimuli in the same class for a given task are marked by black or white text along the axes of the matrices.

A theory relating SUSTAIN’s operation to the brain (11, 22, 23) hypothesizes that HPC forms and alters cluster representations. This notion is similar to computational models of episodic memory that link HPC computations to forming conjunctions of experiences (12, 24). Prefrontal cortex (PFC) is proposed to tune selective attention to features (25–27), as well as direct encoding and retrieval of HPC cluster representations (9, 28–31). In particular, PFC monitors the similarity between the current stimulus information and existing conceptual knowledge and biases HPC functions in reorganizing clusters to reflect goal-relevant features. In other words, what is attended to by PFC affects what is activated in HPC, and how HPC representations are updated impacts how PFC-based attention is tuned. Here, we used the computational model to index each participant’s attentional strategies and organization of object representations across the two learning tasks. We then used these model-based predictions to test how neural representations in HPC for the same experiences dynamically evolve in the face of changing concepts.

An important aspect of our approach is that model-based predictions about dynamic changes in object representations were tailored to each participant’s learning behavior. Using a model-based representational similarity analysis (RSA) approach, for each participant, we compared the similarity structure of the model-predicted cluster representations (i.e., conceptual knowledge) to the neural activation patterns elicited by the insect stimuli (see Fig. 3). We hypothesized that the organization of HPC object representations during learning would track how the model dynamically updated its attention-weighted object representations across the learning tasks. The theorized neural mechanism for such dynamic HPC updating relies on communication between HPC and brain regions important for evaluating sensory and internal mnemonic information (11). Thus, we also predicted a functional coupling between HPC and PFC, subregions of which have been implicated in the formation of generalized knowledge (31, 32) and cognitive control (33).

**Figure 3:**
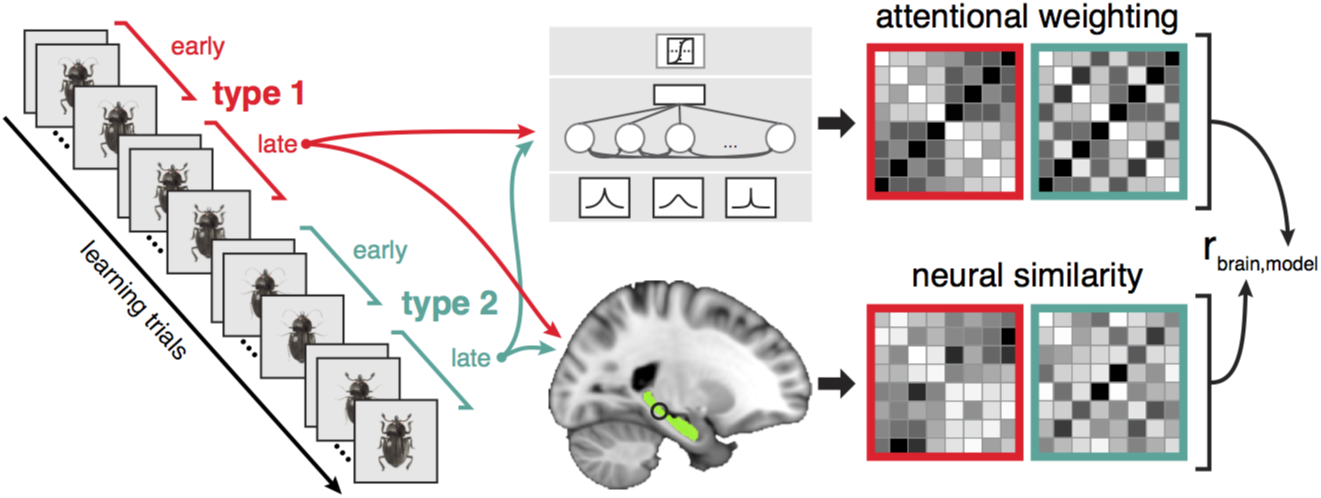
Schematic of model-based representational similarity analysis (RSA). Model predictions and neural measures of stimulus similarity were extracted from the second half of both tasks. For each participant, the learning model was fit to behavior. These optimized model fits were used to generate the representational similarity spaces (Fig. 2c). A searchlight method was used to generate corresponding neural similarity matrices within the hippocampus (highlighted in green) by correlating voxel activation patterns within each searchlight sphere (3 voxel radius) for stimulus pairs from fMRI data recorded during the latter half of the task. The correspondence between model and neural similarity matrices across both tasks was assessed with Spearman correlation.

## Results

### Dynamic updating changes attentional strategies and object similarity

Participants successfully learned both classification problems across learning trial repetitions (Fig. 1c; *β*_*rep*_=0.431, SE=0.046, *z*=9.398, *p*<1×10^−16^) with performance on type 1 reaching asymptote sooner than type 2 (*β*_*task*_=0.928, SE=0.363, *z*=2.556, *p*=0.011). Type 2 learning was relatively slower for participants that learned type 1 first (*β*_*task*order*_=−1.218, SE=0.551, *z*=−2.209, *p*=0.027). No other group level effects on performance reached significance. The learning model was fit separately to each participant’s learning curves and the attention weight parameters (*λ*) were extracted at the end of learning for both tasks (Fig. 2b). According to model predictions, participants allocated attention to the features that were most diagnostic for the given learning task. For the type 1 task, attention was allocated more to the diagnostic dimension *λ*_1_ than the other two dimensions (*Z*s>4.40, *p*s<8×10^−6^). For the type 2 task, attention was allocated more to the two diagnostic dimensions *λ*_2_ and *λ*_3_ (*Z*s>5.80, *p*s<6×10^−9^). This behavioral pattern replicates previous findings (1, 20, 21) and allows us to quantitatively index attention’s influence on HPC conceptual coding.

We also examined the object representations as predicted by the learning model after the concepts had been acquired. For each participant, we extracted the model-based cluster representations for the same stimuli in both learning tasks, operationalized as a vector of values representing the degree that each model cluster was activated by the stimuli. We then calculated the pairwise correlations between these cluster representations (Fig. 2c). Across the two tasks, the similarity structure differed strikingly, reflecting the change in relevancy for the stimulus features; for instance, some stimuli that were less similar in the type 1 problem were more similar in the type 2 problem (e.g., stimuli 1 and 5). This difference in similarity structure across the tasks was confirmed with a randomization test of the matrices’ exchangeability (*Z*=3.42, *p*=0.0024). Moreover, not all stimuli within a category show the same level of similarity (e.g., in type 2, stimuli 1 and 4 are predicted to be very dissimilar despite belonging to the same category). Thus, any neural representations that are found to be consistent with this structure cannot be due simply to the association between stimuli and a category response. Collectively, these behavioral and modeling findings suggest participants learned the tasks by attending to diagnostic information and updating object representations to reflect the distinct attentional strategies required by each task. The similarity structure reflecting model-based object representations were used to test how conceptual coding in HPC dynamically reflected changing task concepts.

### Hippocampal representations change dynamically with model predictions

To evaluate the dynamic nature of HPC-based representations across learning tasks, we measured model-brain consistency with model-based RSA (34). This approach (Fig. 3) allowed us to index the degree that the similarity structure of neural activation patterns matched model-based predictions of conceptual organization. Specifically, we calculated neural similarity between HPC activation patterns for each stimulus pair after the concepts were established in both tasks (i.e., the second half of each task when participants had reached asymptotic performance). The resulting neural similarity matrices, one for each of the two learning tasks, were concatenated and compared to the SUSTAIN similarity matrices with Spearman correlation and a randomization testing procedure. Using searchlight methods (35), this entire process was repeated for all spheres of neural activity (3 voxel radius) within HPC.

The group-level analysis of the model-based RSA (Fig. 4a) revealed a cluster in left anterior HPC (voxelwise threshold *p*<0.005, small volume cluster correction *p*<0.05; cluster peak Z=3.21; cluster peak location: x=−25, y=−15, z=−17; 161 cluster extent) that exhibited significant consistency with the conceptual representations as predicted by the learning model. To visualize the conceptual organization within this HPC region, we derived attention weight estimates from neural similarity measures and projected these weights into stimulus feature space (Fig. 4b). These spaces reflect the influence of attentional tuning with changing task demands; whereas neural representations demonstrated more attention allocated to the first feature dimension (*λ*^n^_1_) in the type 1 task, attention was tuned to the other two feature dimensions (*λ*^n^_2_ and *λ*^n^_3_) in the type 2 task. This HPC region did not vary in response magnitude across tasks (*Z*=0.092, *p*=0.927); all task related modulation was at the level of latent representation. An additional control analysis demonstrated that HPC representational coding was not simply category-based, but rather that attention-weighting inherent to the model was critical to isolate updating mechanisms within HPC (see Supporting Information).

**Figure 4:**
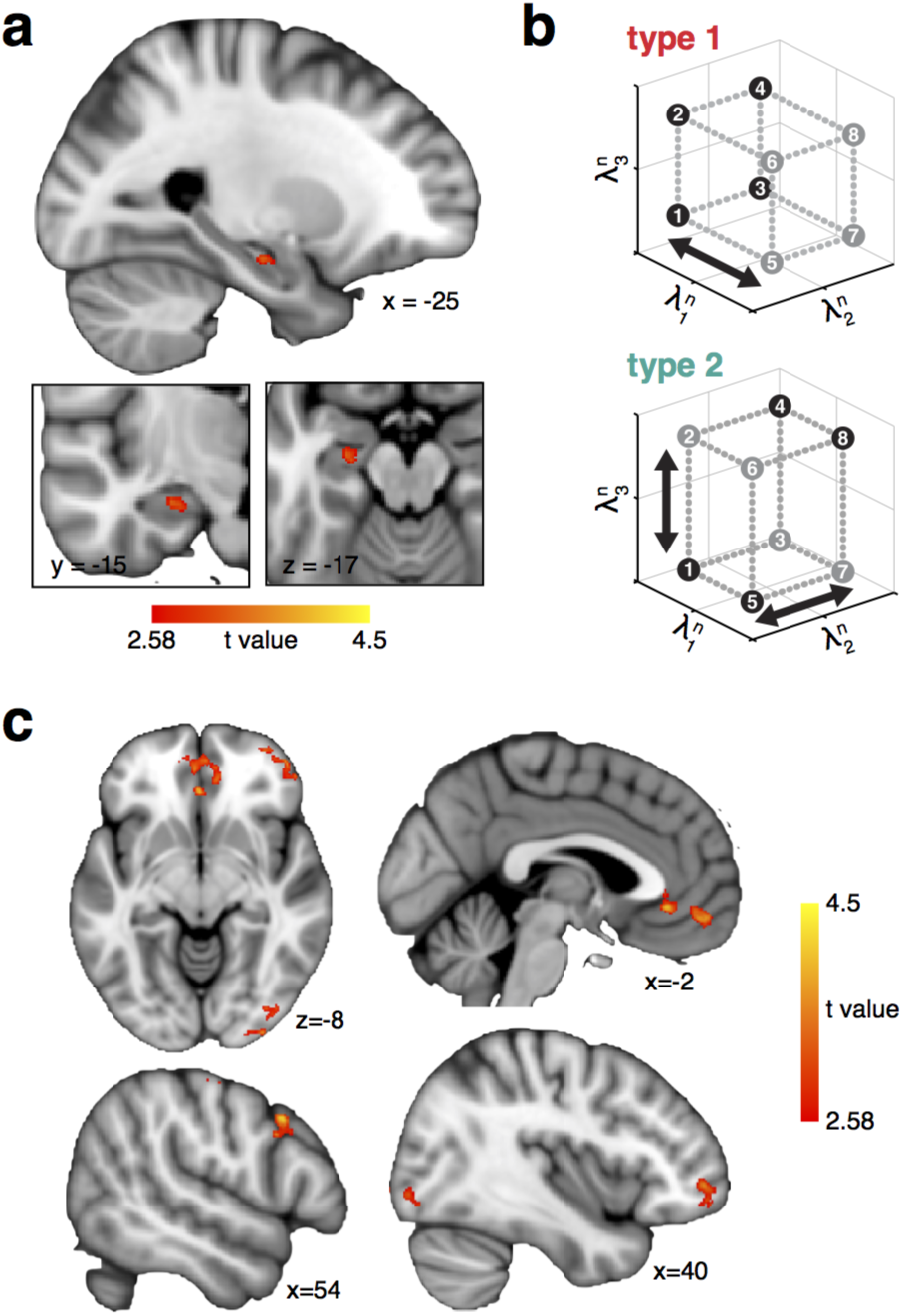
Model-based RSA and learning-related connectivity results. **a)** Neural representations in left anterior HPC were consistent with model predictions of attention-weighted conceptual coding (cluster peak x=−25, y=−15, z=17, 161 voxel cluster extent; voxelwise thresholded at *p*<0.005 and small volume corrected at *p*<0.05 for HPC). **b)** Stimulus-specific neural representations from the HPC region in panel **a** were used to estimate attention weights to the three feature dimensions. These neurally-derived attention weights were then projected into feature space to demonstrate the attentional tuning across tasks. Each point represents a stimulus and is colored according to the class membership for the task. The attention-weighted spaces are a visual depiction of the model-based RSA results (i.e., they are not an independent analysis) and show how attention is tuned across tasks to reconfigure stimulus space into task-relevant conceptual space. The type 1 space suggests attention was allocated more to the diagnostic first dimension (*λ*^n^_1_); the type 2 space reflects a change in attentional tuning, in which attention was allocated more to the other dimensions (*λ*^n^_2_ and *λ*^n^_3_) that were relevant for the task. c) Regions in PFC and occipital cortex showed significantly greater functional coupling with the HPC region identified by model-based RSA during early versus late learning (voxelwise thresholded at *p*<0.005, whole brain cluster extent corrected at *p*<0.05).

### Hippocampal-prefrontal functional connectivity greater during updating

We next evaluated the hypothesis that dynamic updating of HPC representations is facilitated by interactions with PFC (11). We predicted that such interactions would be critical early in learning, when the need for dynamic updating of the conceptual space is most prevalent and the learning model establishes goal-relevant clusters. Specifically, we performed a whole-brain functional connectivity analysis to test whether neural activity in left anterior HPC (Fig. 4a) was coupled with PFC more so during early relative to late learning. Group-level analyses of functional connectivity revealed that both PFC and occipital regions showed enhanced coupling with the HPC seed region in the early learning phase relative to later in learning (Fig. 4c; Table S2). Specifically, activation time courses in bilateral medial prefrontal (mPFC), right frontopolar (FPC), and right dorsolateral prefrontal (dlPFC) cortex were coupled with early learning-related HPC BOLD activity.

## Discussion

Using a model-based fMRI approach, we show that HPC object representations are updated as new concepts are acquired; the same object is represented differently when concepts shift to emphasize new object features. When task demands change, HPC representations are updated to reflect new concepts and when such dynamic updating is occurring, HPC is distinctly coupled with PFC. Furthermore, our approach goes beyond current model-based fMRI methods that examine only the relationship between brain response and individual model parameters. Specifically, we assessed the organization of neural representations and how they change as function of experience through the lens of a computational model and an a *priori* theory linking model to brain regions. By doing so, our approach links formal psychological theory to the neural dynamics of learning (11).

The current findings provide unique support for the hypothesized role of the HPC in building conceptual knowledge (6, 12, 24). Notably, the HPC region showing attention-weighted object representations was predominantly localized to the dentate gyrus/CA_2,3_ region. The intrinsic properties of this region (36, 37) makes it ideal for integrating goal-relevant features into concept representations (6). Although animal (38) and human (39) work has shown support for HPC involvement in the binding of coarse event elements such as items in context (40, 41), the current findings implicate HPC coding at the level of individual stimuli and how they are conceptually organized. Recent work has shed light on the organization of over-learned conceptual representations of visual objects (7, 34, 42); here we show that such conceptual organization can evolve as a function of changing goals. Specifically, by leveraging quantitative model predictions of how attention selects stimulus features impacts the similarity relations among object representations during learning, we demonstrated that HPC coding was sensitive to the stimulus features that were informative to the task at hand.

Two recent human fMRI studies (16, 17) have demonstrated that HPC representations, as evidenced in voxel activation patterns, are distinct for different task states. In these studies, searching through room images for a particular style of wall art evoked distinct HPC patterns relative to searching the same room images for a particular room layout. Although these findings offer compelling evidence that attention enhances encoding of distinct HPC representations, the current study extends beyond this work to characterize *how* that modulation occurs and to show that attention influences the neural representation of learned concepts. We demonstrated that goal-diagnostic information is preferentially encoded into HPC representations, with concept organization evolving as goals change. These results, possible only by linking model predictions to neural representations, provide a substantial contribution towards understanding the computational mechanisms that underlie HPC knowledge formation and updating.

Our findings also add to a growing body of literature suggesting that HPC supports cognitive tasks beyond the domain of episodic memory (43). The finding that anterior HPC forms concept-specific representations speaks to the debate on HPC’s role in representing complex visual objects (44) and classification learning (4, 45). Although findings from seminal rodent and patient studies suggest perirhinal cortex rather than HPC is critical for processing objects composed of multiple features (46–48), the current findings are consistent with the account that HPC is important for organizing complex object representations according to changing contexts (39, 41, 49). Indeed, the current study extends prior fMRI work implicating HPC processes in classification learning (23, 50–52) to demonstrate the influence of goal-relevant selective attention in HPC-based conceptual representations. These results add support for the theoretical proposal from the episodic memory literature that the HPC adaptively builds knowledge representations across episodes (11, 12). It is noteworthy that our findings were localized to the anterior portion of the HPC. Recent rodent (38) and human (8, 53) evidence suggests episodic memories are encoded according to a gradient of generalization along the HPC anterior-posterior axis. Posterior HPC has been shown to exhibit distinct representations for individual episodes, whereas anterior HPC codes for integrated representations that generalize across related episodes (8, 23, 38, 53). The current study builds on these observations to show that anterior HPC representations are not limited to spatial contexts (19, 38, 54, 55) or overlap between related episodes (8), but can also integrate attention-weighted object information across extended learning experiences.

Our connectivity findings speculatively support the view that PFC interacts with HPC when concepts are updated to reflect changes in the learning context. In particular, left anterior HPC showed greater functional connectivity during the early phase of learning with regions of mPFC, a finding predicted by SUSTAIN’s neural framework (11). These results are also consistent with episodic memory findings that PFC biases encoding and retrieval of mnemonic information in HPC (9, 28, 29) and a recent proposal (30) that memories for individual experiences are updated through HPC-mPFC interactions to create generalized knowledge that supports complex behaviors like inference (8, 31). We also found that early-learning HPC activation was coupled with regions implicated in a neural hierarchy of cognitive control along the rostral-caudal axis of lateral PFC (27, 33, 56, 57). Speculatively, such coupling may reflect processes that map stimulus features to goal-specific response and context representations. Finally, that HPC showed coupling with occipital cortex is consistent with previous reports that neural representations in high-level visual areas change as a function of experience (34, 58). Collectively, these findings speculatively offer a potential learning network that should be a target of future studies.

There are relevant parallels between the current study and the extensive literature on task switching (26, 59). This literature seeks to characterize the mechanisms underlying cognitive control in switching between tasks with distinct attentional (60, 61) and response (27, 62) strategies. Here, we tested how individuals engage similar mechanisms of selective attention and control, but within a learning context. By focusing on how participants learned to selectively attend to the most relevant features across changing tasks, we provide a formal account of how attention guides the formation and updating of neural representations of concepts.

In conclusion, the model-based fMRI approach of the current work provides a compelling demonstration that HPC-based object representations are dynamically updated through attention biases. This approach is a unique contribution to the expanding field of computational model-based fMRI methods (63–65). In particular, the model-based RSA method we propose is a fundamental departure from the typical model-based fMRI approach (22, 66, 67) that focuses on localizing time-varying model parameters to activation timeseries of brain regions. Instead, we leverage the structure of the conceptual representations predicted by a learning model to reveal how neural representations of goal-specific concepts are dynamically updated during learning. Furthermore, by marrying computational modeling and neural measures through the shared currency of similarity structure, we leverage RSA in manner that was an original proposed goal of the approach (68), but has seem only limited empirical support (34, 42). With this multifaceted approach, the current study demonstrates that, when goals change, object representations in HPC can be organized in new ways, resulting in updated concepts that highlight the features most critical to the new goal.

## Experimental Procedures

### Subjects and procedures

Twenty-three volunteers participated in the study; all subjects were right handed, had normal or corrected-to-normal vision, and were compensated $75 for participating. After consent in accordance with the University of Texas Institutional Review Board, participants performed all tasks in a 3T Siemens Skyra MRI scanner. They were instructed to learn to classify insect stimuli according to different rules that were based on the combination of the insects’ three features. They were instructed to learn by using the feedback displayed on each trial. The rules that defined the classification problems were not included in any of the instructions; rather, participants had to learn these rules through trial and error. Participants first performed a familiarization task that was included to familiarize participants with the insect stimuli and task procedures to eliminate any neural activation due to stimulus and task novelty during the learning tasks. Participants then performed the type 1 and type 2 classification tasks. For the type 1 task, class associations were defined by a rule depending on the value of one dimension. For the type 2 task, class associations were defined by an XOR logical rule that depended on the value of the two dimensions that were not relevant in the type 1 task. The order of the type 1 and 2 tasks was counterbalanced across participants. The classification tasks consisted of learning trials presented in an event-related design. On each trial, an insect image was presented for 3.5s and participants made a response as to the insect’s class. The stimulus presentation period was followed by a 0.5-4.5s fixation, a 2s feedback screen consisting of the insect image, text of whether the response was correct or incorrect, and the correct class, and then a 4-8s fixation. Each of the eight insect images was presented in four learning trials during each fMRI run and participants completed four fMRI runs for each classification task. Whole brain fMRI data was acquired with 1.7mm isotropic voxels and a TR of 2 seconds. Full procedures and MRI data acquisition and processing details are described in Supplemental Information Methods.

### Model-based representational similarity analysis

We fit SUSTAIN, a computational learning model, to each participant’s learning performance. Stimuli were presented to SUSTAIN in the same order as what the participants experienced and model parameters were optimized to predict each participant’s learning performance in the familiarization task and two learning tasks. The optimized parameters were used to extract measures of dimensional attention weights and latent representations (model cluster activation vectors) of the stimuli during the second half of learning in the two tasks. The pairwise similarities of the cluster activation vectors were then calculated with Pearson correlation to generate similarity matrices. Values from the upper triangle of the task-specific similarity matrices were concatenated and these served as the model-based prediction of attention-biased representations. Neural similarity matrices for the stimuli were calculated by first estimating activation patterns for each stimulus using an event-specific univariate general linear model (GLM) approach (69). The neural similarity between stimulus-specific activation patterns from the second half of learning was assessed with a searchlight method such that the Pearson correlation was calculated for all pairwise stimulus-specific activation patterns from a searchlight sphere with a radius of 3 voxels. Values from the upper triangle of the task-specific neural similarity matrices were concatenated to serve as the neural similarity between the stimuli across tasks. The correspondence between the resulting neural similarity matrix and model-based similarity matrix was then assessed with Spearman correlation and a reshuffling randomization test. This searchlight method was applied to all searchlight spheres within HPC. For more details, see Supplemental Information Methods, Computational modeling analysis and Model-based RSA.

### Functional connectivity analysis

Left anterior HPC functional connectivity with the rest of the brain was assessed using voxelwise regression. Mean activation time courses from the left anterior HPC region identified in the model-based RSA were extracted for each participant and entered into a GLM with the time course as a regressor. The resulting parameter estimates from the first two runs (early learning) were contrasted with the last two runs (late learning). The resulting contrast images were normalized to MNI space and submitted to group analysis. For more details, see Supplemental Information Methods, Functional connectivity analysis.

## Author Contributions

M.L.M., B.C.L., and A.R.P. designed research; M.L.M. performed research; M.L.M. analyzed data; and M.L.M., B.C.L., and A.R.P. wrote the paper.

## Acknowledgments

We thank M. Schlichting, B. Gelman, E. Stein, and K. Nguyen for help with data collection. Thank you also to the Texas Advanced Computing Center (http://www.tacc.utexas.edu) at The University of Texas at Austin for critical computing resources. M.L.M. is supported by the National Institute of Mental Health (F32-MH100904). A.R.P. is supported by the National Science Foundation (CAREER Award 1056019), and the National Institute of Mental Health (R01-MH100121). B.C.L is supported by the Leverhulme Trust (RPG-2014-075), a Wellcome Trust Senior Investigator Award (WT106931MA), and the National Institute of Child Health and Human Development (1P01HD080679).

